# Modeling pulsed evolution and time-independent variation improves the confidence level of ancestral and hidden state predictions in continuous traits

**DOI:** 10.1101/2021.03.29.437517

**Authors:** Yingnan Gao, Martin Wu

## Abstract

Ancestral state reconstruction is a fundamental tool for studying trait evolution. It is also very useful for predicting the unknown trait values (hidden states) of extant species. A well-known problem in ancestral and hidden state predictions is that the uncertainty associated with predictions can be so large that predictions themselves are of little use. Therefore, for meaningful interpretation of predicted traits and hypothesis testing, it is prudent to accurately assess the uncertainty of the predictions. Commonly used Brownian motion (BM) model fails to capture the complexity of tempo and mode of trait evolution in nature, making predictions under the BM model vulnerable to lack-of-fit errors from model misspecification. Using simulations and empirical data (bacterial genomic traits and vertebrate body size), we show that the presence of pulsed evolution and time-independent variation significantly undermines the confidence level of continuous traits predicted under the BM model. The residual Z-scores are neither homoscedastic nor normally distributed. Consequently, the 95% confidence intervals of predicted traits are so unreliable that the actual coverage probability ranges from 29% (strongly permissive) to 100% (strongly conservative). To remedy the model misspecification problem, we develop RasperGade that accounts for both pulsed evolution and time-independent variation. When applied to simulated and empirical data, RasperGade outperforms commonly used tools such as *ape*. It restores the normality and homoscedasticity of the Z-score distributions. Accordingly, RasperGade greatly improves the reliability of confidence intervals of predictions and reduces the deviation of their actual coverage probabilities from the 95% expectation by as much as 99%.

## Introduction

Ancestral state reconstruction of continuous traits is important in comparative biology and the study of trait evolution, as in most cases the traits of ancient species are very difficult, if ever possible, to measure. For instance, to study the origin of life, it would be useful to know the optimum growth temperature (*T*_*OPT*_) of the last universal common ancestor (LUCA), which is impossible to measure directly. However, *T*_*OPT*_ can be inferred from the GC content of the ribosomal RNA (rRNA) gene, which in turn can be estimated from the rRNA genes of extant species through ancestral state reconstruction (Galtier et al. 1999). Ancestral state reconstruction of continuous traits also lays the foundation of many phylogenetic measures that include the phylogenetically independent contrast (Felsenstein 1985) and phylogenetic signal (Blomberg and Garland 2002). Beyond the traits of ancient species, ancestral state reconstruction can also be applied to infer the unknown traits of extant species, in an approach known as the hidden state prediction (Garland and Ives 2000; Zaneveld and Thurber 2014). Hidden state prediction is especially useful when traits of extant species are difficult to measure. For example, the vast majority of environmental bacterial species have not been cultured but their functional traits can be inferred by hidden state prediction using the molecular phylogeny of 16S rRNA gene (Kembel et al. 2012; Langille et al. 2013).

For continuous traits, model-based methods are widely used to reconstruct the ancestral states (Webster and Purvis 2002; Royer-Carenzi and Didier 2016). The commonly used Brownian motion (BM) model assumes that trait is under neutral selection, and that small incremental changes accumulate at a constant rate over time or the branches in the phylogeny. It follows that trait evolves gradually with the amount of evolution accumulating in proportion to the branch length, and the trait change along a branch follows a normal distribution (Felsenstein 1985; Schluter et al. 1997). Felsenstein provided an analytic solution to the maximum-likelihood estimate (MLE) of the root ancestral state under the BM model (Felsenstein 1985; Maddison 1991), which does not require the time-consuming optimization procedure. In addition to the MLE of the root ancestral state, Felsenstein’s method also provides the standard error of the estimated root ancestral state (Felsenstein 1985), which can be used to assess the uncertainty of the ancestral state prediction. Under the BM model and other time-reversible models, hidden state prediction can be simply done by rerooting the tree using the unknown tip as the root and then reconstructing the new root’s ancestral state (Garland and Ives 2000).

One well-known challenge to ancestral state prediction is the inherent uncertainty associated with the prediction, which increases with increasing evolutionary distance between the nodes to be predicted and observed nodes (Oakley and Cunningham 2000). The uncertainty of the predicted ancestral state is often so large that the predictions are of little use, especially for the distant ancestors (Schluter et al. 1997; Webster and Purvis 2002). Taking *T*_*OPT*_ as an example, a predicted *T*_*OPT*_ may have a confidence interval so wide that it spans the full range of bacterial optimum growth temperature (from psychrophilic to thermophilic), making the prediction meaningless in practice. Therefore, in addition to getting an accurate point estimate of ancestral or hidden state, it is equally important to obtain a reliable confidence measure of the prediction. As explained below, model misspecification can produce misleading confidence intervals of the predictions.

Although the BM model is convenient for quick ancestral and hidden state predictions, it has many limitations imposed by its assumptions. In nature, most traits are rarely under truly neutral selection, violating the first assumption of the BM model. Although a study has shown that fluctuating directional selection can be seen as neutral in the long term (Hansen and Martins 1996), on shorter time scales, diversifying and purifying selections can increase and decrease the trait evolution rate respectively and therefore the constant evolution rate assumption of the BM model is also frequently violated in nature. Studies of fossil records and comparative traits have shown that trait evolution can proceed in rapid bursts separated by long periods of stasis (Simpson 1944; Eldredge and Gould 1972; Jackson and Cheetham 1994). Such pulsed evolution has been shown to be prevalent in animals (Uyeda et al. 2011; Landis and Schraiber 2017). Modeling pulsed trait evolution using the BM model results in model misspecification, which in turn could lead to erroneous ancestral state reconstruction. When trait evolves by pulsed evolution, the normality of trait changes, especially of those along short branches, breaks and consequently the Gaussian likelihood surface no longer holds. It follows that predictions of ancestral and hidden states based on the normal distribution, both the point estimate and the confidence interval, may be inaccurate. To address this problem, stable distribution, a heavy-tailed analog to the normal distribution, has been applied to ancestral state reconstruction (Elliot and Mooers 2014). Although it can capture the variation in the trait evolution rate, the lack of a closed-form probability function and the need of using a Bayesian approach due to the rugged likelihood surface have limited its use in large phylogenies.

One often neglected problem for ancestral and hidden state predictions is the presence of time-independent variation. As its name suggests, time-independent variation is not explained by phylogeny. There are several sources of time-independent variation, which includes the measurement error of the trait (Landis and Schraiber 2017), the heritability of the trait (Lynch 1991), and short-lived variation and lineages (Futuyma 2010; Rosenblum et al. 2012). Accounting for time-independent variation is important for reliably predicting the ancestral and hidden states, as increased time-independent variation has been shown to increase the mean error of ancestral state reconstruction as it erodes the phylogenetic signal (Litsios and Salamin 2012). Time-independent variation also contributes to the inherent uncertainty.

The lack of known ancestral states (e.g., fossil records) has always been an obstacle in evaluating the performance of ancestral state reconstruction methods. With a few exceptions, such evaluation has been mostly limited to simulations (Royer-Carenzi and Didier 2016). For reconstruction methods that use time-reversible models (e.g., the BM model, the stable model and the pulsed evolution model), leave-one-out cross validating the hidden state prediction can overcome this obstacle (Kembel et al. 2012), because predicting the hidden states is essentially the same as reconstructing the ancestral states over a rerooted phylogeny under time-reversible models (Garland and Ives 2000; Kembel et al. 2012; Zaneveld and Thurber 2014).

Here we developed RasperGade, a method of ancestral and hidden state prediction that accounts for both time-independent variation and pulsed evolution by extending Felsenstein’s recursive reconstruction method. Using simulated data, we show that RasperGade improves the accuracy and the confidence level of the predicted ancestral and hidden states compared to the commonly used methods that assume the constant-rate BM model. Using empirical data and leave-one-out cross validation, we demonstrate that RasperGade outperforms the BM-based methods in hidden state prediction by greatly improving the confidence level. Our results suggest that, when predicting the ancestral and hidden states of continuous traits, the tempo and mode of evolution should always be assessed and taken into account.

## New Approaches

Commonly used BM model-based ancestral and hidden state prediction methods such as those implemented in the R package *ape* fail to capture pulsed evolution and time-independent variation. To address this model misspecification problem, we introduce a new algorithm that incorporates pulsed evolution and time-independent variation into ancestral state reconstruction. Under the BM model, the trait change between any two nodes is normally distributed, and consequently in reconstruction, the ancestral state is also normally distributed (Felsenstein 1985; Schluter et al. 1997) and its maximum likelihood estimate can be calculated by the algorithm introduced by Felsenstein (Felsenstein 1985; Maddison 1991). This algorithm can be easily extended to incorporate time-independent variation, by adding a positive variance to tip trait values, wherein the normality of trait changes and ancestral states is maintained. However, when pulsed evolution is present, trait changes along branches no longer follow a normal distribution as the stochastic nature of jumps introduces uncertainty into the average rate, and neither does the ancestral state. To remedy the model misspecification problem caused by pulsed evolution, we can decompose the non-normal distribution of the trait changes into a weighted mixture of normal distributions, and apply Felsenstein’s method of reconstruction to each component in the mixture. Specifically, because we model pulsed evolution as jumps that occur at a constant rate and whose magnitudes follow a normal distribution, the trait change introduced by *n* jumps along a branch is normally distributed. Consequently, we can decompose the non-normal distribution of trait change on each branch by the number of jumps that might have occurred along the branch. For ancestral state reconstruction, we can sum up the distributions of all possible combinations of jumps along the two descendent branches using Felsenstein’s method. The weighted mixture of these distributions will be the actual probability distribution of the ancestral state in the presence of pulsed evolution, and instead of calculating its MLE, we use the mean of the distribution as the point estimate. Following this strategy, we implemented RasperGade (Reconstructing Ancestral State under Pulsed Evolution in R by Gaussian Decomposition), a set of R functions that predict ancestral and hidden states while accounting for pulsed evolution and time-independent variation. More details of RasperGade are available in the Materials and Methods and Supplementary Text sections.

## Result

### Pulsed evolution and time-independent variation are present in bacterial genomic trait evolution

Previous studies revealed that pulsed evolution and time-independent variation are prevalent in vertebrate body size evolution (Uyeda et al. 2011; Landis and Schraiber 2017), leading us to speculate that they might also be present in bacterial trait evolution. To test our hypothesis, we determined four traits (ribosomal RNA GC%, genomic GC%, genome size, and the average nitrogen atoms per residual side chain) from 6,668 complete bacterial genomes. These traits have been linked to bacterial fitness (Wang et al. 2006; Lauro et al. 2009; Martinez-Gutierrez and Aylward 2019). Following Landis and Schraiber (Landis and Schraiber 2017), we formulate three different evolution models: the classic Brownian motion model (BM) for gradual evolution, Brownian motion plus time-independent variation model (BM+ε), and Brownian motion plus pulsed evolution and time-independent variation (BM+PE+ε). We model pulsed evolution using the Levy process with normal jumps as described in (Landis and Schraiber 2017). Briefly, we assume that jumps occur at a constant rate along the branches, and thus the number of jumps on a certain branch follows a Poisson distribution. The trait change introduced by one jump follows a normal distribution with a mean of zero and a constant variance. We model time-independent variation by a normally distributed random variable. We implemented a maximum-likelihood framework to estimate the parameters of the three models from the phylogenetically independent contrasts and compared their ability to explain the trait variation in extant bacterial genomes through AIC. We found that the BM+PE+∊ model provides the best fit to all bacterial traits (Table S1), with its AIC weight being greater than 99.9% in all cases. Specifically in the BM+PE+ε model, we found that pulsed evolution and the time-independent variation contribute up to 95% and 0.9% of the trait variation respectively.

### Pulsed evolution and time-independent variation induce deviant Z-score distributions

The presence of pulsed evolution and time-independent variation violates the assumption of constant trait evolution rate in the BM model, and consequently it may undermine the reliability of the predicted traits and their predicted uncertainty. To test our hypothesis, we first carried out a simulation study. To establish a baseline for evaluating the reliability of the predictions (both the point estimate and its uncertainty), we simulated and predicted traits both using the BM model. Under the BM model, the residual error of each predicted ancestral or hidden state is expected to be normally distributed. It follows that the normalized residual Z-score should follow a standard normal distribution and should be homoscedastic over the predicted standard error. When pooling the ancestral or hidden states of nodes in a phylogeny together, dependence among nodes will cause small deviations from the above expectation, but the overall normality and homoscedasticity should be maintained. We found that this is the case in our simulation (Figures 1 and S1, A and B) and we will refer to it as the “BM baseline” thereafter. We use two statistics to quantify the normality and heteroscedasticity of the residual Z-score distributions. The D statistic in the Kolmorogov-Smirnov test is used to measure the goodness-of-fit of the Z-scores to the standard normal distribution to assess its normality. The median chi-square (median **χ**^2^) in the Fligner-Killeen test of the Z-scores over predicted standard error is used to measure the degree of its heteroscedasticity. To simplify the interpretation of results, we normalize the D statistics and median **χ**^2^ of the “BM baseline” to 1 (Table 1 and Table S2).

**Figure 1.**
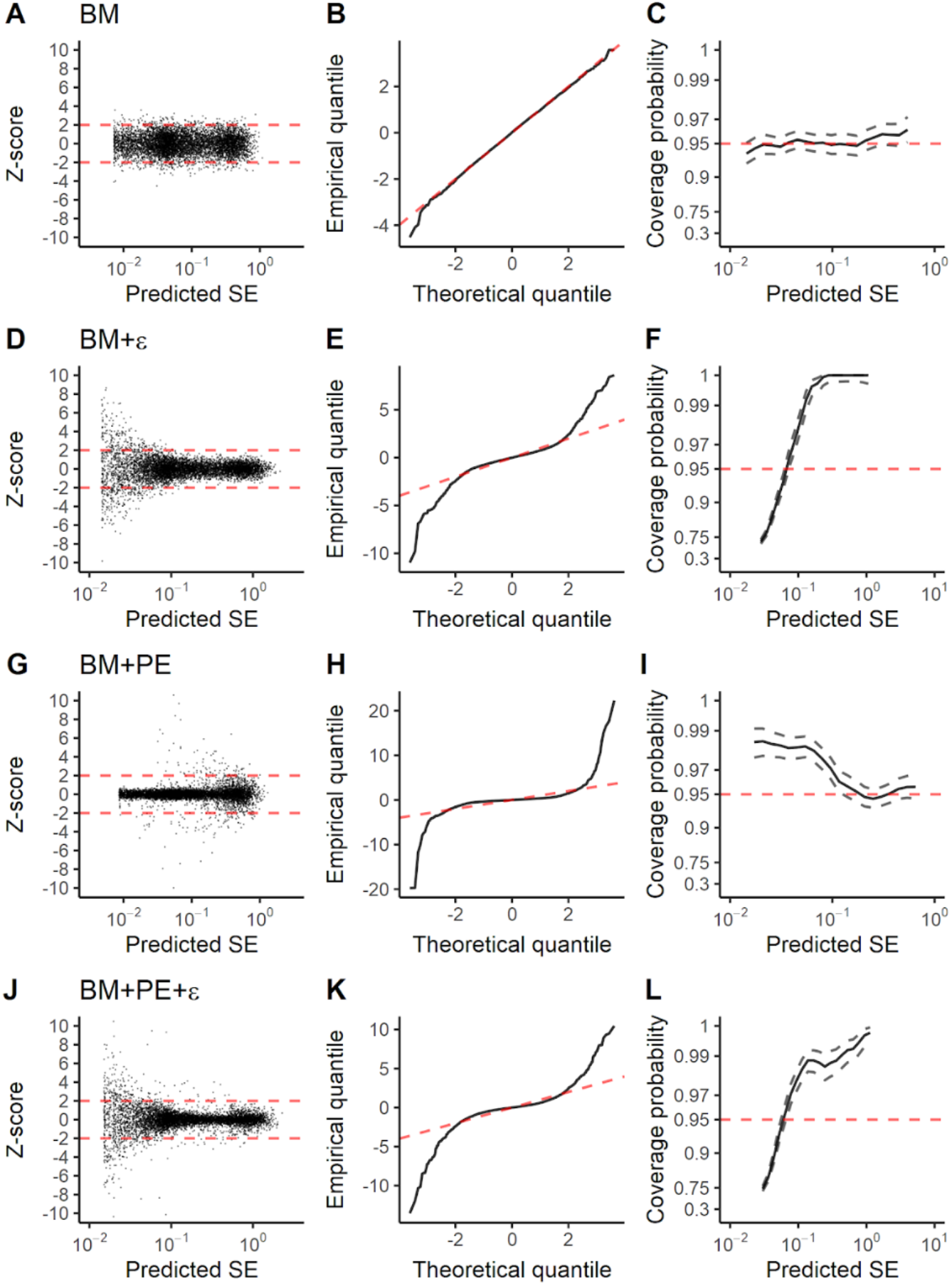
Diagnostic plots of residual errors in hidden state prediction using the BM model on data simulated with various evolutionary models. The first to the last row shows results from data simulated with the BM, BM+ε, BM+PE and BM+PE+ε model respectively. The first column (A, D, G and J): the distribution of residual Z-scores along the predicted standard error. The red dashed lines represent the expected 95% confidence interval when the Z-scores follow the standard normal distribution. The second column (B, E, H and K): the Q-Q plot of the distribution of the residual Z-scores against the standard normal distribution. The dashed red line shows the expectation when the two distributions are identical. The third column (C, F, I, and L): the actual coverage probability of the 95% confidence interval plotted along the predicted standard error. The red dashed line represents the expected 95% coverage probability, and the grey dashed lines delimit the 95% confidence interval for the actual coverage probability. The axis of the coverage probability is transformed as described in the Materials and Methods section.

**Table 1.** The performance of hidden state prediction in leave-one-out cross-validation using the BM model and the true models.

| Model used in simulation | Model used in hidden state prediction | Performance statistics |  |  |  |  |
| --- | --- | --- | --- | --- | --- | --- |
| | | Error | Bias | D | Median $\chi^2$ | RSS<sub>CP</sub> |
| BM | BM | $1.222 \times 10^{-1}$ | $7.94 \times 10^{-5}$ | 1.00 | 1.00 | 1.00 |
| BM+$\epsilon$ | BM | $1.374 \times 10^{-1}$ | $2.10 \times 10^{-4}$ | 8.66*** | 99.11*** | 36.76*** |
| | BM+$\epsilon$ | $1.367 \times 10^{-1}$*** | $2.08 \times 10^{-4}$ | 1.00 | 1.02 | 0.94 |
| BM+PE | BM | $9.823 \times 10^{-2}$ | $-2.38 \times 10^{-4}$ | 22.01*** | 38.48*** | 15.87*** |
| | BM+PE | $9.467 \times 10^{-2}$*** | $-1.86 \times 10^{-4}$ | 2.12*** | 1.42*** | 1.09 |
| BM+PE+$\epsilon$ | BM | $1.208 \times 10^{-1}$ | $-4.88 \times 10^{-5}$ | 14.09*** | 106.81*** | 25.66*** |
| | BM+PE+$\epsilon$ | $1.163 \times 10^{-1}$*** | $-1.01 \times 10^{-4}$ | 1.32*** | 1.11* | 0.96 |

Note: The table presents the performance statistics of hidden state prediction in leave-one-out cross-validation by the BM and the true models. For each simulation, we ran 100 replicates and present the average statistics. The D, median $\chi^2$ and RSS<sub>CP</sub> statistics are normalized by their BM baselines (the first row). Error: the mean absolute difference between the predicted and the true trait value. Bias: the mean difference between the predicted and the true trait value. D: the test statistic in Kolmogorov-Smirnov test of residual Z-scores to the standard normal distribution. Median $\chi^2$ : the test statistic in Fligner-Killeen test of residual Z-scores' homoscedasticity. RSS<sub>CP</sub>: the sum of squares of the actual coverage probabilities of the 95% confidence intervals to the 95% expectation. We tested if the mean of bias is different from zero, if the D, median $\chi^2$ and RSS<sub>CP</sub> predicted using the true models are greater than those of the BM baselines (normalized to 1), and if the mean of error when predicting with the true model is different from that when predicting with the BM model. The significance levels are shown by the asterisks: \*\*\* <0.001, \*\* 0.001-0.01, \* 0.01-0.05.

Once the baseline is established, we then simulated trait evolution with pulsed evolution and time-independent variation, in which pulsed evolution and time-independent variation contribute 90% and 0.5% of the trait variation on average, respectively. We then predicted traits using the BM model. For the point estimate of ancestral and hidden states, we found that regardless of the model used to simulate the trait evolution, the bias of prediction is not significantly different from zero (P≥0.096, Table 1 and Table S2). The error of the prediction, on the other hand, varies from 0.06 to 0.14 (Table 1 and Table S1), corresponding to 6% to 14% change in traits such as genome size or body size. However, when pulsed evolution or time-independent variation is present, it is no longer appropriate to use the BM model to assess the uncertainty of the predictions: the residual Z-scores are neither homoscedastic (Figures 1 and S1, D, G, and J) nor normally distributed (Figures 1 and S1, E, H and K). Specifically, the median **χ**^2^ that measures the degree of heteroscedasticity increases up to 107-fold compared to the median **χ**^2^ of the BM baseline (P<0.001 Table 1 and Table S2), and the D statistic that measures the normality is inflated up to 22-fold compared to the BM baseline (P<0.001, Table 1 and Table S2). Consequently, the uncertainty predicted using the BM model is unreliable. In addition, we found that the deviation from the normal distribution and the degree of heteroscedasticity increase as the relative contribution of pulsed evolution and magnitude of time-independent variation increase (Figure S2, first and second columns).

### Pulsed evolution and time-independent variation produce misleading confidence intervals

The predicted uncertainty of an ancestral or hidden state describes how the true trait value distributes around the predicted value, and thus it can be applied to generate confidence intervals for hypothesis testing. Because model misspecification results in deviant Z-score distributions, we expect that it also undermines the reliability of confidence intervals. When we examined the 95% confidence interval of each prediction, we found that the actual coverage probability, the empirical frequency that the confidence interval covers the true trait value, deviates from the 95% expectation when pulsed evolution or time-independent variation is present (Figures 1 and S1, F, I and L). When time-independent variation is present, the 95% confidence interval tends to be strongly permissive (having lower actual coverage probability than the 95% expectation) when the predicted standard errors are small, and can reach as low as 70.53% for predicted hidden states and 29.38% for ancestral states. The 95% confidence interval becomes conservative (having higher actual coverage probability than the 95% expectation) as the predicted standard error increases, and can reach 100.00% in both ancestral and hidden state predictions. Compared to the BM baseline, the residual sum of squares between the actual coverage probability and the expected 95% (RSS_CP_) inflates up to 37-fold and 23-fold on average in hidden and ancestral state predictions respectively. When pulsed evolution is present, the 95% confidence interval tends to be conservative for small or intermediate predicted standard errors, reaching 98.56% for predicted hidden states and 98.65% for ancestral states. But it does not appear to be permissive for any predicted standard error. In this case, the RSS_CP_ inflates 16-fold and 7-fold on average in hidden and ancestral state predictions respectively. We found that the RSS_CP_ increases when the contribution of pulsed evolution and time-independent variation increase (Figure S2, the third column).

### The BM model produces unreliable confidence estimates of hidden states in the empirical data

Because pulsed evolution and time-independent variation contribute substantially to bacterial and vertebrate trait variation, we speculate that deviant distributions of residual errors similar to those observed in simulations are also present in empirical data. Since we did not know the ancestral states of these traits, we tested our hypothesis by leave-one-out cross-validation in hidden state predictions. As expected, similar deviant patterns were observed in all bacterial traits and in vertebrate body size. Residual Z-scores show strong heteroscedasticity over the predicted standard error (Figure 2, the first column), and the median **χ**^2^ inflates from 12-fold to 82-fold compared to the BM baseline (Table 2). On the other hand, residual Z-scores show drastic deviation from the standard normal distribution (Figure 2, the second column), as the D statistic inflates from 14-fold to 22-fold compared to the BM baseline (Table 2). We also observed that the actual coverage probability of the 95% confidence interval deviates substantially from the expected 95% (Figure 2, the third column), with the RSS_CP_ inflates from 14-fold to 29-fold compared to the BM baseline. For rRNA and genomic GC%, the 95% confidence interval changes from being normal to being conservative and permissive when the predicted standard error increases, with the minimum and maximum of the coverage probability reaching 85.24% and 99.70% respectively. For genome size, N-ARSC and body size, the 95% confidence interval changes from being overly permissive to being overly conservative, with the minimum coverage probability dropping to 75.67% and the maximum coverage probability reaching 100.00%.

**Figure 2.**
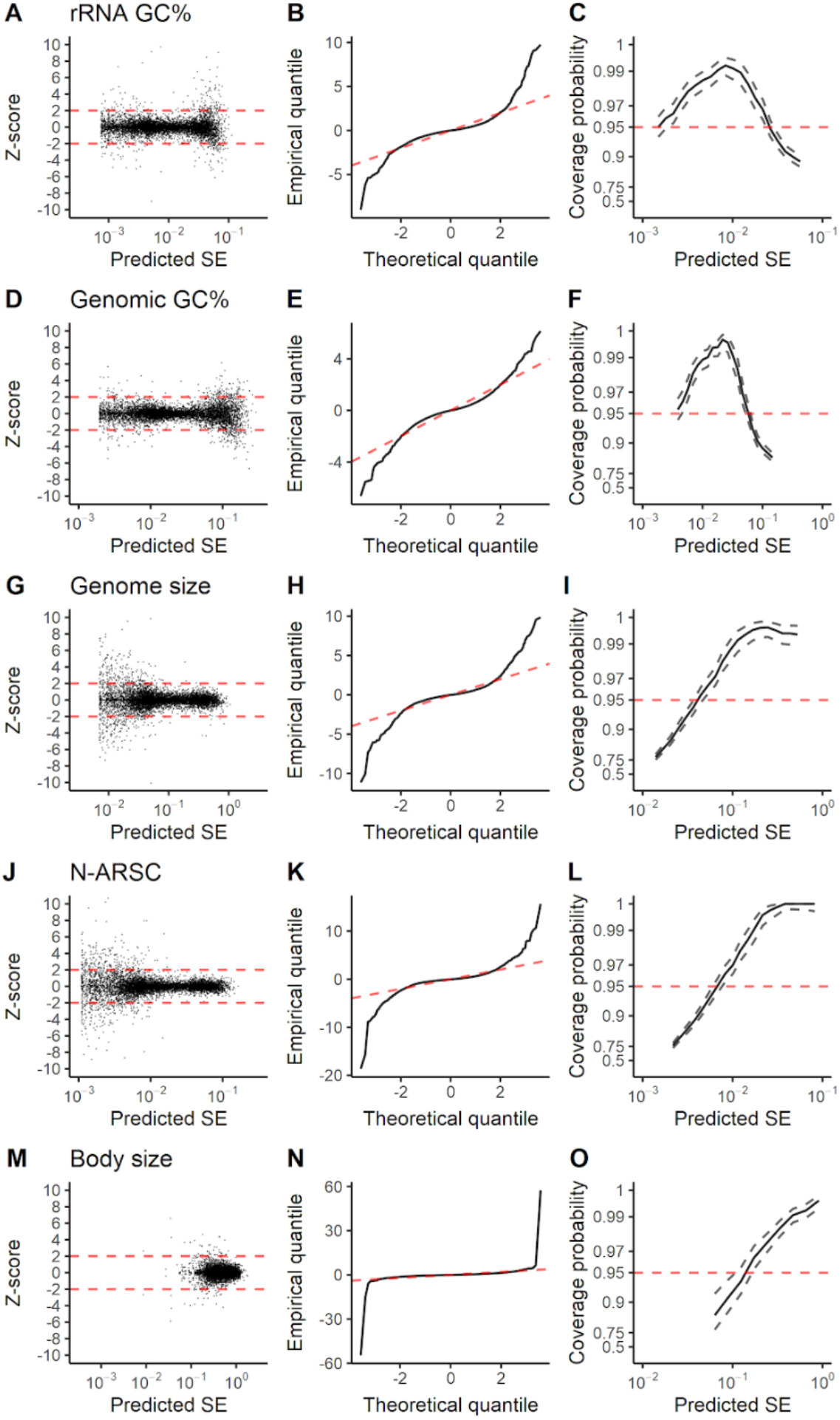
Diagnostic plots of residual errors in hidden state prediction using the BM model on bacterial and vertebrate trait data. The first to the last row shows the results of rRNA GC%, genomic GC%, genome size, N-ARSC and vertebrate body size respectively. The column layout is the same as that of Figure 1.

**Table 2.**
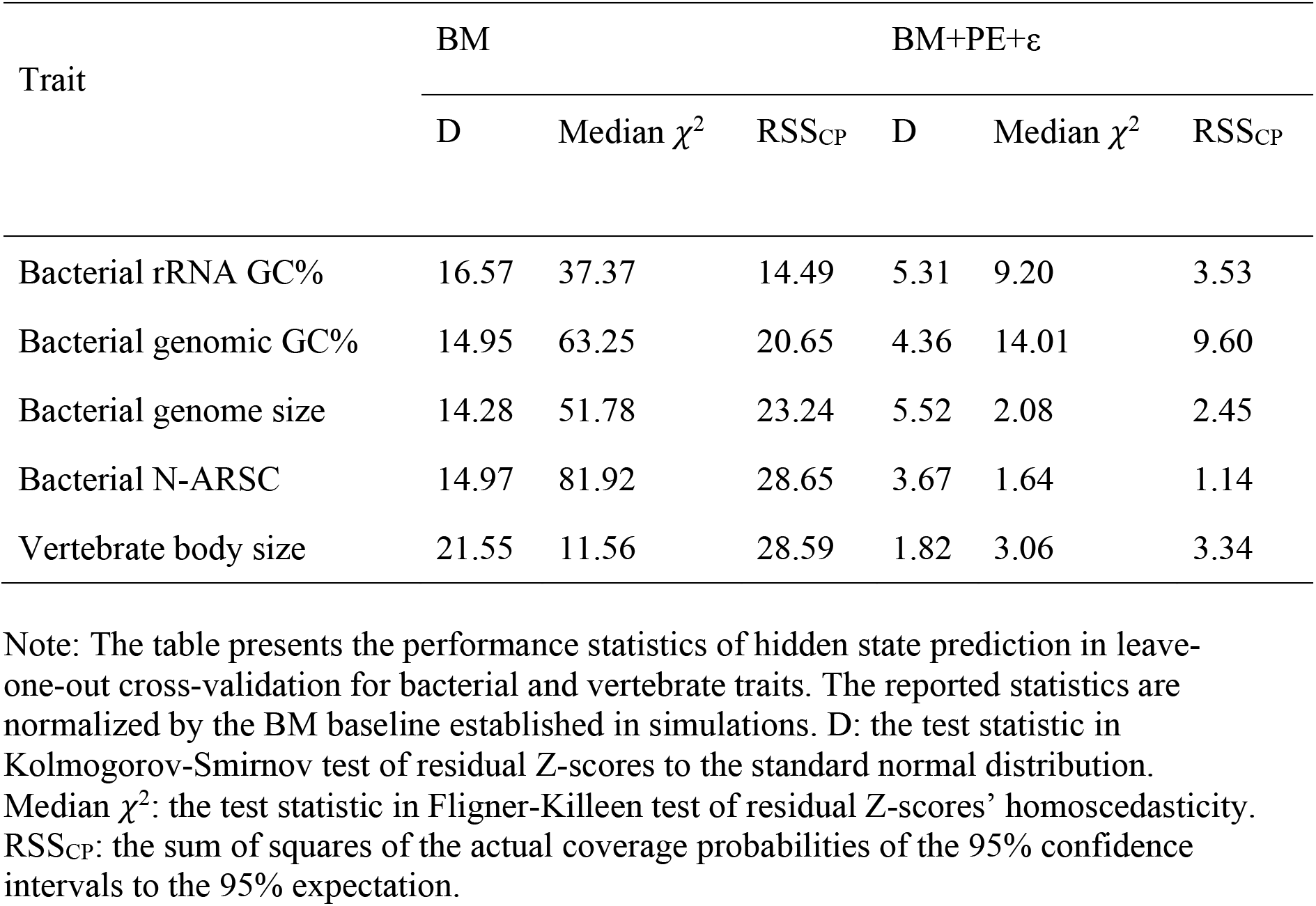
Performance of hidden state prediction in bacterial and vertebrate traits using the BM and BM+PE+ε models.

| Trait | BM | | | BM+PE+$\epsilon$ | | |
| --- | --- | --- | --- | --- | --- | --- |
| | D | Median $\chi^2$ | RSS<sub>CP</sub> | D | Median $\chi^2$ | RSS<sub>CP</sub> |
| Bacterial rRNA GC% | 16.57 | 37.37 | 14.49 | 5.31 | 9.20 | 3.53 |
| Bacterial genomic GC% | 14.95 | 63.25 | 20.65 | 4.36 | 14.01 | 9.60 |
| Bacterial genome size | 14.28 | 51.78 | 23.24 | 5.52 | 2.08 | 2.45 |
| Bacterial N-ARSC | 14.97 | 81.92 | 28.65 | 3.67 | 1.64 | 1.14 |
| Vertebrate body size | 21.55 | 11.56 | 28.59 | 1.82 | 3.06 | 3.34 |

Note: The table presents the performance statistics of hidden state prediction in leave-one-out cross-validation for bacterial and vertebrate traits. The reported statistics are normalized by the BM baseline established in simulations. D: the test statistic in Kolmogorov-Smirnov test of residual Z-scores to the standard normal distribution. Median $\chi^2$ : the test statistic in Fligner-Killeen test of residual Z-scores' homoscedasticity. RSS<sub>CP</sub>: the sum of squares of the actual coverage probabilities of the 95% confidence intervals to the 95% expectation.

### RasperGade restores the normality and homoscedasticity of pseudo-Z-scores

Because the confidence interval of the predictions made under the BM model is greatly undermined by the presence of pulsed evolution and time-independent variation, we developed RasperGade to remedy the model misspecification problem. First we tested if applying RasperGade improves the outcome of ancestral and hidden state predictions in simulated data. We found that RasperGade yields marginal but significant reduction in the error of the predictions (Table 1 and Table S2, P<0.001), and the predictions remain unbiased (Table 1 and Table S2, P≥0.085). Despite the small improvement in accuracy, RasperGade greatly reduces the heteroscedasticity in the pseudo-Z-scores, an analog of Z-scores for non-normal residual errors, over the predicted standard error (Figures 3 and S3, the first column). The inflation in median **χ**^2^ drops by more than 98% (P<0.001). In addition, RasperGade restores the standard normal distribution of pseudo-Z-scores (Figures 3 and S3, the second column) and reduces the inflation in the D statistic by more than 94% (Table 1 and Table S2, P<0.001). The actual coverage probabilities become much closer to the expected 95% than predictions made using the BM-based method, ranging from 93.43% to 96.44% (Figures 3 and S3, the third column), and the inflation in RSS_CP_ drops by more than 98% and they are no longer significantly different from the BM baseline (P≥0.051). The performance of RasperGade is not sensitive to the relative contribution of pulsed evolution and to the magnitude of time-independent variation (Figure S2). This suggests that RasperGade is robust as long as the model specified is sufficiently accurate.

**Figure 3.**
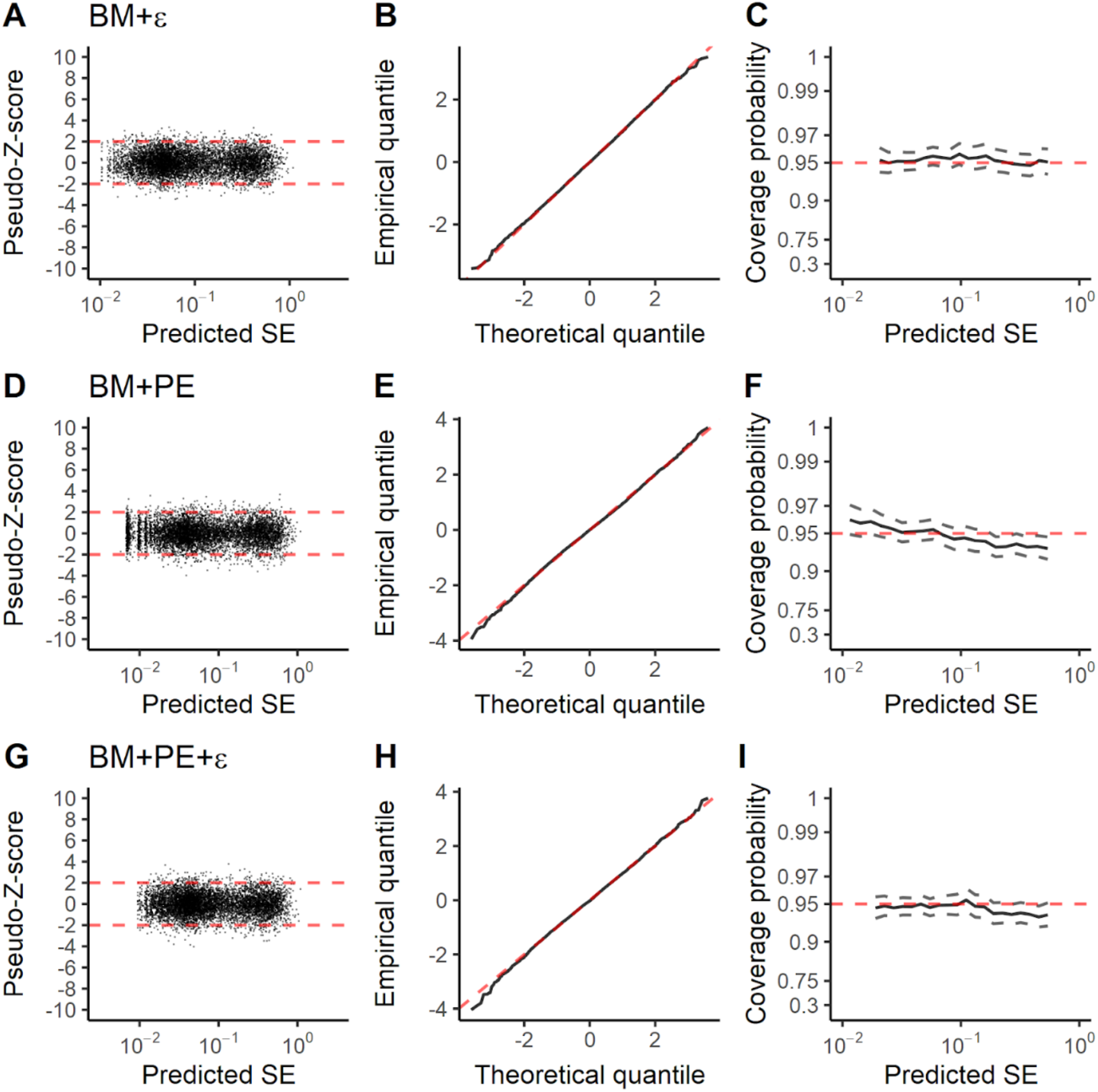
Diagnostic plots of residual errors in hidden state prediction by RasperGade using the true model on data simulated with various evolutionary models. The first to the last row shows results from data simulated with the BM+ε, BM+PE and BM+PE+ε model respectively. The column layout is the same as that of Figure 1.

### RasperGade improves the uncertainty assessment in empirical data

Because RasperGade greatly improved our ability to assess the uncertainty of predictions in simulation, we applied it to the bacterial and vertebrate traits and evaluated its performance. We found that although RasperGade does not eliminate heteroscedasticity, it does significantly reduce the degree of heteroscedasticity in pseudo-Z-score (Figure 4, the first column), with the inflation of median χ^2^ decreasing by 77% to 99% (Table 2). In addition, applying RasperGade partially restores the normality of the pseudo-Z-score distribution (Figure 4, the second column), with the inflation of D statistic reduced by 66% to 96% (Table 2). Compared to methods using the BM model, RasperGade greatly improves the actual coverage probability of the 95% confidence interval (Figure 4, the third column), with the inflation in RSS_CP_ decreased by 56% to 99%. For rRNA GC%, genome size, N-ARSC and body size, the actual coverage probability ranges from 90.92% to 96.64%. For genomic GC%, the actual coverage probability ranges from 89.68% to 98.95%.

**Figure 4.**
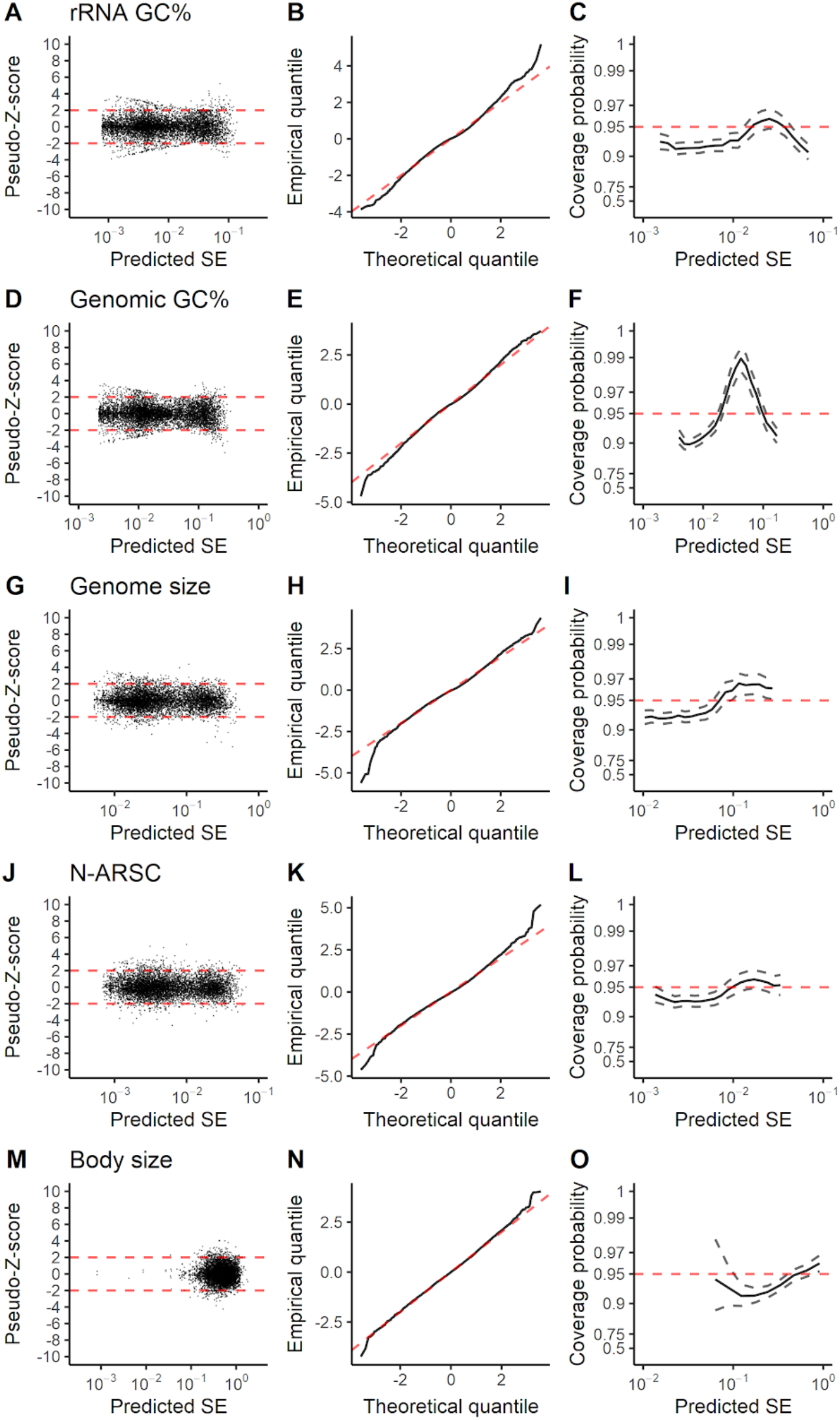
Diagnostic plots of residual errors in hidden state prediction by RasperGade using the BM+PE+ε model on bacterial and vertebrate trait data. The layout is the same as that of Figure 2.

### Extending RasperGade to discrete traits

Strictly speaking, the BM model of trait evolution only applies to continuous traits. However, it has been extended to discrete traits such as the 16S rRNA gene copy number (GCN) whose values are consecutive integers (Kembel et al. 2012; Langille et al. 2013). Similarly, we can extend the pulsed evolution model and RasperGade to this type of discrete traits. To avoid excessive zero-contrasts, we first add a normally distributed random number with a mean of zero and a small variance (jittering variance) to each trait value (which will be subtracted from the estimated variance of time-independent variation at the end), and predict traits as if they are truly continuous. We round the point estimate of the ancestral or hidden state to the nearest integer, and integrate the probability density of nearby trait values to represent the probability of the predicted integer trait value. For example, if the point estimate of the ancestral state is 1.1, RasperGade will predict that the ancestral state is 1 (nearest integer). It will calculate the probability of the true trait value being 1 by integrating the probability of the trait value between 0.5 and 1.5. We use residual sum of squares (RSS) between the empirical frequency of correct predictions and the predicted probability to evaluate the goodness-of-fit of the predicted uncertainty.

Our model testing shows that pulsed evolution dominates the evolution of 16S rRNA GCN, explaining close to 100% of the trait variation. On the other hand, time-independent variation contributes less than 0.2% of the variation. Therefore, we used 16S rRNA GCN to evaluate the performance of RasperGade. When cross validated by hidden state prediction, we found that compared to methods that use the BM model, RasperGade reduces the RSS from 49.84 to 11.22, greatly improving the reliability of the predicted uncertainty. In practice, we can use the predicted probability to classify the predicted trait value as either reliable if the predicted probability is greater than 95%, or as unreliable otherwise. Such classification can be applied to select a set of high-quality predictions for downstream analyses. Using the probability predicted under the BM model results in a total classification error rate (correct predictions classified as unreliable or incorrect predictions classified as reliable) of 69.80%, while using the probability predicted by RasperGade using the BM+PE+∊ model, the error rate reduces to only 33.43%.

## Discussion

The errors in ancestral and hidden state predictions come from two major sources: the inherent uncertainty of the trait evolution, and the lack-of-fit errors introduced by model misspecification. The inherent uncertainty roots from the trait evolution’s stochastic nature, and it increases with the evolutionary distance between the to-be-predicted node and the observed nodes (Oakley and Cunningham 2000). As a result, the inherent uncertainty cannot be reduced unless additional observations (e.g., fossil records or increased taxon sampling) are introduced (Maddison 1995; Martins 1999; Salisbury and Kim 2001; Finarelli and Flynn 2006). Nevertheless, the inherent uncertainty is predictable from the tempo and mode of trait evolution, and the predicted uncertainty can and should be used to guide the interpretation and application of predicted ancestral and hidden states. Although the inherent uncertainty in ancestral state reconstruction can be so high that the predictions are no longer meaningful (especially for deep nodes) (Webster and Purvis 2002), it is usually much smaller in hidden state predictions. This is because the hidden states often have closely related sister tips with known states (Martins 1999).

The lack-of-fit errors, on the other hand, are caused by model misspecification. The BM model, widely used for its simplicity, fails to capture the complexity of the tempo and mode of trait evolution in nature, making predictions under the BM model vulnerable to lack-of-fit errors from model misspecification. Using simulation and empirical data, we show that pulsed evolution and time-independent variation can cause serious problems for commonly used methods and tools that assume the BM model. In simulation, when pulsed evolution and/or time-independent variation are present, the residual Z-scores for predictions made using the BM model are neither normal nor homoscedastic. As a result, the 95% confidence intervals of the predicted trait values are so unreliable that the actual coverage probability varies from 29% to 100%. Such deviation greatly undermines the confidence levels of the predicted traits.

Because pulsed evolution follows a compound Poisson process, the rate of evolution, although relatively constant on a longer time scale, varies greatly on a shorter time scale. Consequently, the variation introduced by pulsed evolution along short branches is not proportional to the branch length. When using the BM model for prediction, the model misspecification caused by pulse evolution will be most severe for short branches and will be minimal for long branches, as manifested in the degree of deviation from the expectation in both the Z-scores and the coverage probability of the 95% confidence intervals (Figure 1 G and I).

Time-independent variation has many sources. The bacterial traits in this study are based on high-quality genomic sequences, and thus the time-independent variation in these traits is unlikely due to either low heritability or measurement errors. A more likely source of time-independent variation in these bacterial traits is the ephemeral variation that does not persist over time. Because time-independent variation does not accumulate over time, its relative contribution to the trait variation diminishes as the branch length increases and the variation from the BM and pulsed evolution accumulates. As a result, time-independent variation has the largest impact on the Z-scores of predictions with short branches (Figure 1 D). Additionally, time-independent variation strongly interferes with the fitting of BM model, as the fitted model will try to compensate for the time-independent variation in the short branches with an inflated rate of trait evolution, which in turn will have a large impact on the actual coverage probability of 95% confidence intervals for both short and long branches (Figure 1 F), even though the contribution of time-independent variation to the total trait variation is small and a strong phylogenetic signal is maintained.

Unlike the inherent uncertainty, the lack-of-fit errors can be reduced or eliminated by using a more realistic model of trait evolution. RasperGade is designed to rectify the two types of model misspecification discussed above by incorporating pulsed evolution and time-independent variation. RasperGade marginally but significantly improves the point estimates of ancestral or hidden states. However, the true strength of RasperGade lies in its ability to greatly improve the 95% confidence levels of the predictions. It reduces the deviation of the actual coverage probability of the confidence intervals from the 95% expectation by as much as 99%. It does so by restoring the normality and homoscedasticity of the Z-score distributions. Such improvement makes interpretation of the predicted trait values and downstream analysis much more meaningful and reliable.

RasperGade does likelihood-based model selection among three models (BM, BM+ε, BM+PE+ε) and selects the best model for trait predictions. The predictions of RasperGade using the BM model are identical to those made using the *ace* function of the R package *ape* (with Felsenstein’s method). RasperGade’s performance is robust to the varying degrees of pulsed evolution and time-independent variation (Figure S2). However, like any model-based methods, RasperGade’s performance depends on accurate model parameters. In practice, we recommend that a phylogeny with at least 200 tips should be used to get a reliable estimation of the model parameters.

As the vast majority of bacterial species have not been cultured and very little is known about their biology and phenotypic traits, one obvious RasperGade application is to predict unknown traits of these uncultured bacterial species based on their positions in the extremely well sampled 16S rRNA phylogeny. For example, the 16S rRNA GCN has been correlated with the life strategy of the bacterial species, with oligotrophs having fewer copies of 16S rRNA gene than copiotrophs (Lauro et al. 2009; Roller et al. 2016). The 16S rRNA GCN is also needed for correctly estimating the composition (relative species abundance) of a microbial community using 16S rRNA sequencing data (Kembel et al. 2012). However, it has been suggested that the uncertainty associated with 16S rRNA GCN prediction varies greatly from taxon to taxon (Louca et al. 2018). Therefore, keeping high quality predictions and removing the bad ones is an essential quality control step for high quality downstream analysis. In this study, we show that RasperGade greatly improves the confidence level of the 16S rRNA GCN prediction by reducing the overall classification error rate from 69.80% to 33.43%. Using a similar approach, PICRUST predicts the gene copy numbers of metabolic genes and thus the metabolic capacity of an uncharacterized bacterial species or a microbial community from 16S rRNA sequences (Langille et al. 2013). Currently PICRUST uses the BM model for hidden state prediction and for estimating the prediction uncertainty. Based on the results of this study, we speculate that RasperGade will also improve the confidence intervals of PICRUST predictions.

RasperGade has some limitations. For example, RasperGade assumes no evolutionary trend. A persisting evolutionary trend in trait evolution has been shown to reduce the accuracy of the ancestral state reconstruction (Schluter et al. 1997; Garland and Díaz-Uriarte 1999; Oakley and Cunningham 2000; Finarelli and Flynn 2006), and thus it should be handled by its corresponding evolutionary model such as the arithmetic Brownian motion model (Royer-Carenzi and Didier 2016). However, this may be less of a concern for hidden state predictions using an ultrametric tree, as the evolutionary trend cancels each other out when the tree is rerooted using the hidden state tip as the root. To reduce the computational cost, RasperGade uses one normal distribution to approximate the true distribution of marginal ancestral states. As a result, in the presence of pulsed evolution, it does not completely restore the normality and homoscedasticity of the Z-scores even when the true model is used for predictions (Table 1 and Table S1). In empirical data, RasperGade does not fully restore the normality and homoscedasticity of the Z-scores. One possibility is that RasperGade assumes that the tempo and mode of trait evolution is homogeneous throughout the phylogeny, which might not be the case. It is also possible there are more than one type of jumps occurring in bacterial and vertebrate trait evolution. Although not implemented in this study, RasperGade has the potential to handle these complex models with modifications in how to conduct the Gaussian decomposition.

In summary, for ancestral and hidden state predictions of continuous traits, we recommend always first testing the presence of pulsed evolution and/or time-independent variation, and using RasperGade for predictions when either is present.

## Materials and Methods

### Preparation and compilation of trait and phylogenetic data

We examined five bacterial genomic traits and one vertebrate trait in this study. For bacteria, we downloaded 10,616 complete bacterial genomes from the NCBI RefSeq database on September 6, 2018. From each genome we identified 31 “universal” bacterial protein-coding marker genes using AMPHORA2 (Wu and Scott 2012) and constructed a bacterial genome tree based on the concatenated and trimmed protein sequence alignment of the marker genes using FastTree (Price et al. 2010). For better resolution, we re-optimized the branch length of the genome tree with the DNA sequence alignments of the marker genes using RAxML (Stamatakis 2014). We removed genomes with identical alignments, extremely long branches, ambiguous bases or unreliable annotations from the genome trees. For each of the 6,668 bacterial genomes remained, we calculated 4 continuous traits: the ribosomal RNA stem GC% (rRNA GC%), genomic GC% (GC%), genome size (excluding plasmids) and the average nitrogen atoms per residual side chain (N-ARSC), and one discrete trait, the 16S rRNA gene copy number (16S rRNA GCN). We transformed continuous traits (logit-transformation for rRNA GC% and GC%; log-transformation for genome size and N-ARSC) to make them comply with the assumption of continuous trait evolution, and assumed that the evolution of discrete 16S rRNA gene copy number is well approximated by a continuous model without any transformation. For vertebrates, we used phylogenies and the log-transformed body size data of 64 vertebrate clades (excluding Chelonia in reptiles and *Etheostoma* in fish) compiled by Landis and Schraiber (Landis and Schraiber 2017).

### Modeling pulsed evolution and time-independent variation

We model pulsed evolution using the Levy process with normal jumps as described by Uyeda et al. (Uyeda et al. 2011) and Landis and Schraiber (Landis and Schraiber 2017) for its simplicity, and implemented custom R scripts to estimate the parameters of pulsed evolution and time-independent variation using a maximum likelihood framework. Specifically, we model time-independent variation by a normal distribution and estimate its variance ε. We model the occurrence of jumps in pulsed evolution by a Poisson process, and estimate its rate λ and the size of the jumps *M*, which is the variance of trait change introduced per jump. We also estimate *r*, the rate of Brownian motion. For bacterial traits, we fitted the models on the full bacterial phylogeny. For vertebrates, we first fitted the models on each clade separately using the R package *pulsR* (Landis and Schraiber 2017). We then excluded 28 clades of vertebrates in which pulsed evolution is not present and jointly fitted the models on the remaining 36 clades using our custom R scripts.

### Simulating trait evolution with pulsed evolution and time-independent variation

We simulated trait evolution over the bacterial phylogeny with two schemes. In the scheme for case studies, we simulated trait evolution with fixed degrees of pulsed evolution and time-independent variation to evaluate their impact on ancestral and hidden state predictions and RasperGade’s ability to remedy the model misspecification problem. Specifically, we set the magnitude (variance) of time-independent variation to 4×10^−3^ (approximately 0.5% of total trait variation in tip values), the rate of pulsed evolution to 5 jumps per unit branch length, the contribution of pulsed evolution to 90%, and the average rate of trait evolution (average variance introduced in trait values per unit branch length) to 1. We used a full factorial design of the presence of pulsed evolution and time-independent variation and simulated trait evolution with four models: the BM model, the BM+ε model, the BM+PE model, and the BM+PE+ε model. Each model was simulated with 100 replicates.

In the second scheme, we simulated trait evolution with varying degrees of pulsed evolution or time-independent variation to evaluate the trend of their impact on ancestral and hidden state predictions and the robustness of RasperGade. In all simulations, the average rate of trait evolution per unit branch length was set to 1. For time-independent variation, we simulated without pulsed evolution, and set the time-independent variation’s magnitude (variance) to 0, 4×10^−4^ or 4×10^−3^. For pulsed evolution, we simulated without time-independent variation, and set its contribution to 0%, 10%, 30%, 50%, 70%, 90% or 100%, with an average of 5 jumps per unit branch length. Each combination of parameters was simulated with 100 replicates.

### Incorporating pulsed evolution and time-independent variation in ancestral state reconstruction

We establish the method for reconstructing ancestral states with pulsed evolution and time-independent variation by extending Felsenstein’s method under the BM model (Felsenstein 1985). For simplicity, we only illustrate reconstruction over a pair of nodes in a bifurcating tree as shown in Figure S4 A, but the method does not require dichotomy in general. Under the BM model with a fixed rate, the likelihood surface of the ancestral states is a Gaussian (normal) one, and consequently the maximum-likelihood estimate of the ancestral state of an internal node can be calculated analytically:

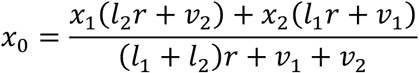

where x_1_ and x_2_ are the trait values of the descendants, l_1_ and l_2_ are the branch lengths to the descendants, r is the rate of BM (gradual trait evolution), and v_1_ and v_2_ are the uncertainty associated with the descendants’ trait values measured as variances. And the variance of the reconstructed ancestral state is:

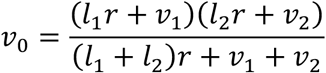

Under the BM model, the uncertainty associated with trait values of tip nodes is usually defaulted to zero, or the squared standard error of the mean if the trait value is averaged from a population. The time-independent variation is incorporated into the ancestral state reconstruction simply by adding its variance (ε) to the uncertainty of the tip nodes.

Compared to time-independent variation, expanding the BM model of ancestral state reconstruction to incorporate pulsed evolution is less straightforward, as the likelihood surface of ancestral states is no longer a simple Gaussian function. To address such complexity, we decompose the likelihood surface of ancestral states by the number of jumps occurring on each branch, so that in each decomposed component, the trait change on each branch is normally distributed. In one decomposed component, given a certain number of jumps on each branch, the conditional ancestral state is normally distributed and its mean is

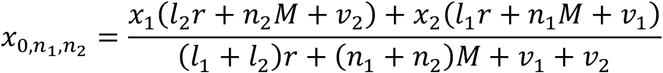

where *n*_1_ and *n*_2_ are the number of jumps occurring on branch *l*_*1*_ and *l*_*2*_, respectively, and *M* is the variance of the trait change introduced per jump. Similarly, the variance of this conditional ancestral state can be derived as

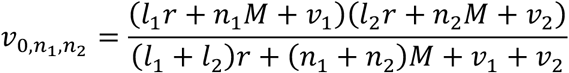

And the probability weight of each conditional ancestral state is

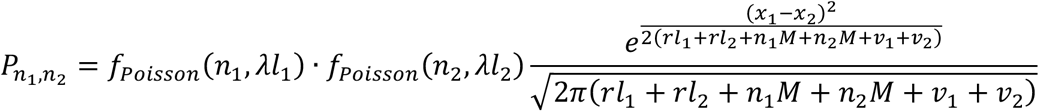

where λ is the frequency of jump per unit branch length, and *f*_*poisson*_ is the Poisson probability mass function for *n* jumps along a branch of length *l*. The likelihood surface of the ancestral state can be derived by mixing these normally distributed conditional ancestral states together. However, the maximum-likelihood estimate for this likelihood surface is not easy to derive, and instead we use the mean of the ancestral state as the point estimate. This mean is given by the weighted mean of all conditional ancestral states:

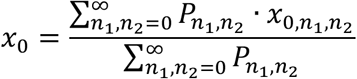

and the variance of the ancestral state is

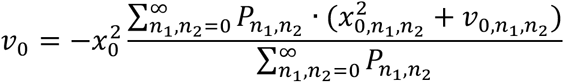

Under the BM model, the uncertainty of predicted ancestral and hidden states can be fully predicted by the variance of the ancestral or hidden states, as the residual errors are normally distributed with a mean of zero and a variance equal to that of the ancestral or hidden states. With time-independent variation, the residual errors are still normally distributed, but the variance of the time-independent variation should be added to the distribution of residual errors. With pulsed evolution, the distribution of residual errors is a mixture of normal distributions with different means and variances. As a result, we have to describe the uncertainty of predicted ancestral and hidden states using the cumulative distribution function:

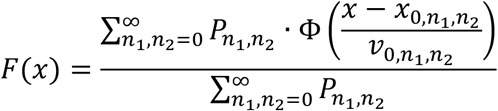

where x is the trait value and Φ(*x*) is the cumulative distribution function for the standard normal distribution.

All the marginal ancestral states and their uncertainty along the phylogeny can be calculated using the above formulas from tips to the root following Felsenstein’s method (Felsenstein 1985) (Figure S4 B). To reduce the time required to calculate all marginal ancestral states, we approximate the uncertainty of marginal ancestral states with a normal distribution with the same mean and variance as the true distribution. Global ancestral states and hidden states can be calculated by rerooting the phylogeny with the focal node or tip as the root and reconstructing the ancestral state at the new root using the above formulas (Kembel et al. 2012) (Figure S4 C). We do not simplify the uncertainty of global ancestral states and hidden states. In this study, all hidden states are predicted by the leave-one-out cross-validation as described by Kembel et al. (Kembel et al. 2012).

### Calculation of pseudo-Z-score

Each ancestral or hidden state has its own uncertainty distribution, and in order to evaluate the performance of RasperGade, we need to normalize the ancestral and hidden states’ residual error so that they follow the same distribution. When residual errors follow normal distributions with different variances, the standard deviations of these normal distributions (the standard errors) are used to normalize the residual errors into the residual Z-scores, which follow the standard normal distribution. When residual errors are not normally distributed, such residual Z-scores can no longer be used. Instead, we find a pseudo-Z-score (Z_pseudo_) for each ancestral or hidden state that satisfy the following equation:

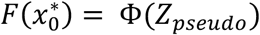

where F(x) is the predicted cumulative distribution of the ancestral or hidden states, Φ(*x*) is the cumulative distribution function for the standard normal distribution, and 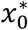 is the true trait value for the ancestral or hidden state. The pseudo-Z-scores will follow the standard normal distribution if the predicted cumulative distribution of the residual error is accurate.

### Evaluating the performance of ancestral and hidden state prediction

Ancestral and hidden state predictions provide two types of predictions: a point estimate of the trait and a predicted distribution describing its associated uncertainty. The accuracy of a point estimate is evaluated by bias and error as defined in Kembel et al. (Kembel et al. 2012), where bias is the mean of the difference between the point estimate and the true value, and error is the mean of the absolute difference between them. For uncertainty, we evaluate the goodness-of-fit between the predictions’ Z-scores and the standard normal distribution, first visually by the Q-Q plot and then statistically by the D statistic in the Kolmogorov-Smirnov test. The degree of heteroscedasticity of the uncertainty is evaluated through the median χ^2^ statistic in Fligner–Killeen test by dividing the Z-scores into 20 sorted groups based on their predicted standard error (SE). The actual coverage probability of the 95% confidence interval over predicted SE is calculated using a sliding window, which spans from 0.5 SE to 2 SE. Within each window, we calculate the actual coverage probability *p* as the percentage that the true trait value lies within the 95% confidence interval of the prediction. When plotting the coverage probability, we transform the coverage probability with the following formula so the deviation between the actual coverage probability and the 95% expectation is evenly distributed around 95%:

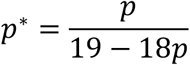

The deviation of actual coverage probability from the 95% expectation is measured by residual sum of squares (RSS_CP_) summed over the predicted SE. Specifically, we divide the Z-scores into 20 sorted groups based on their predicted SE, calculate the actual coverage probability *p* in each group, and calculate the RSS_CP_ from the coverage probabilities in these groups using the formula:

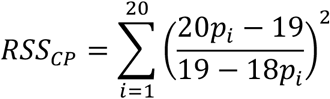

where *i* is the index of each group. For convenient interpretation, we normalize the D statistic, the median **χ**^2^ and the RSS_CP_ by their baseline values from 100 simulations and predictions under the BM model.

### Extending RasperGade to integer trait

Because integer traits exhibit characteristics similar to those of continuous traits, we can extend RasperGade to integer traits by making some modifications to the ancestral and hidden prediction procedures. We fit the evolution model to the integer trait as if it is a truly continuous trait. To avoid excessive contrasts with a value of zero that will interfere with the maximum-likelihood estimation, we add a small white noise with variance of 10^−8^ to the integer trait values and subtract it from the estimated time-independent variation parameter before the reconstruction. In ancestral and hidden state predictions, we round up the predicted ancestral or hidden state to the nearest integer. In terms of the predicted uncertainty, we derive the predicted probability that the predicted value is identical to the true value by integrating the predicted uncertainty distribution. For example, if we predict that the ancestral state is 1, we will derive the probability that the true trait value is 1 by integrating the predicted uncertainty distribution from 0.5 to 1.5. We evaluate its goodness-of-fit by examining if the empirical frequency of correct predictions matches the predicted probability. Specifically, we bin the predicted probability from 0 to 1 into 100 bins, and within each bin, we calculate the average predicted probability 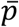 and the empirical frequency *p.* We transform the empirical probability with the formula

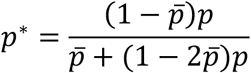

Bins without predictions are omitted. The goodness-of-fit is calculated as

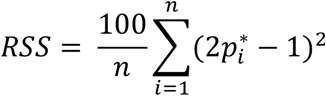

where *n* is the actual number of bins that have at least one prediction. We do not test heteroscedasticity for integer traits.

### Software

RasperGade can be downloaded as a R package from https://github.com/wu-lab-uva/RasperGade.

## Supplementary Text

### Gaussian decomposition of the likelihood surface of ancestral states

Considering a simple but general case as in Figure S4 A, the trait changes over the branches can follow any distribution, and thus the likelihood of the ancestral state x_0_ given the descendant trait values x_1_ and x_2_ can be expressed as:

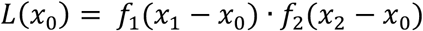

where L(x) denotes the likelihood function and f_1_(x) and f_2_(x) denote the probability density function of trait changes over the two branches. We suppose that f_1_(x) and f_2_(x) can be decomposed into the weighted sum of several normal distributions:

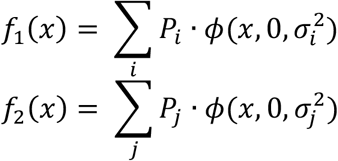

where *ϕ*(*x*, 0, σ^2^) denotes normal probability density functions with a mean of 0 and a variance of *σ*^2^, *P* denotes the weight of each normal distribution, and *i* and *j* denote the indices of these normal density functions. Then the likelihood function L(x_0_) above can be expressed as:

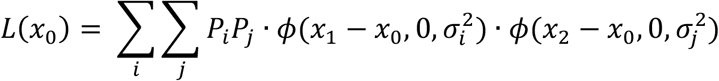

This likelihood function L(x_0_) can be further simplified to the sum of several weighted normal density functions:

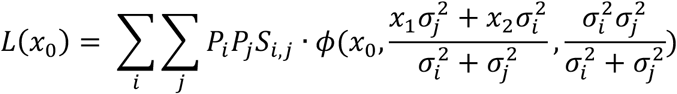

where 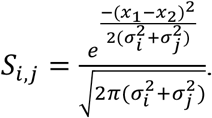

In general, for any distribution, there is no universal analytic solution to how it can be decomposed into a weighted sum of normal distributions. However, under the pulsed evolution model where normal jumps occur at a constant rate, the trait change over the branches can be described by the probability density function:

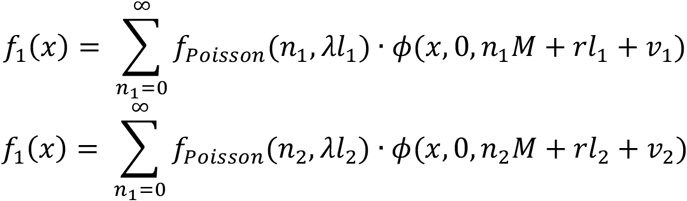

where *f*_*poisson*_ is the Poisson probability mass function for *n* jumps along a branch of length *l*, *r* is the rate of Brownian motion, *v*_1_ and *v*_2_ are the variance of uncertainty associated with trait *x*_1_ and *x*_2_, respectively, λ is the frequency of jump per unit branch length, *M* is the variance of jump size, and n_1_ and n_2_ are the number of jumps occurring on each branch. These probability density functions fit the Gaussian decomposition equations described above by setting *i* = *n*_1_, *j* = *n*_2_, 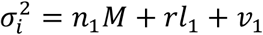, 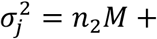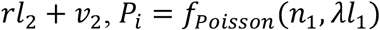 and *P*_*Poisson*_. Consequently, the likelihood function L(x_0_) under the pulsed evolution model can fit the Gaussian decomposition equation by setting

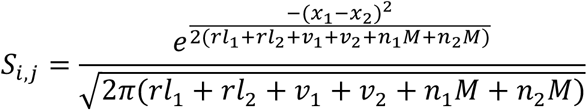

As a result, Gaussian decomposition can be done under the pulsed evolution model through decomposing by the number of jumps occurring on each branch.

**Figure S1.**
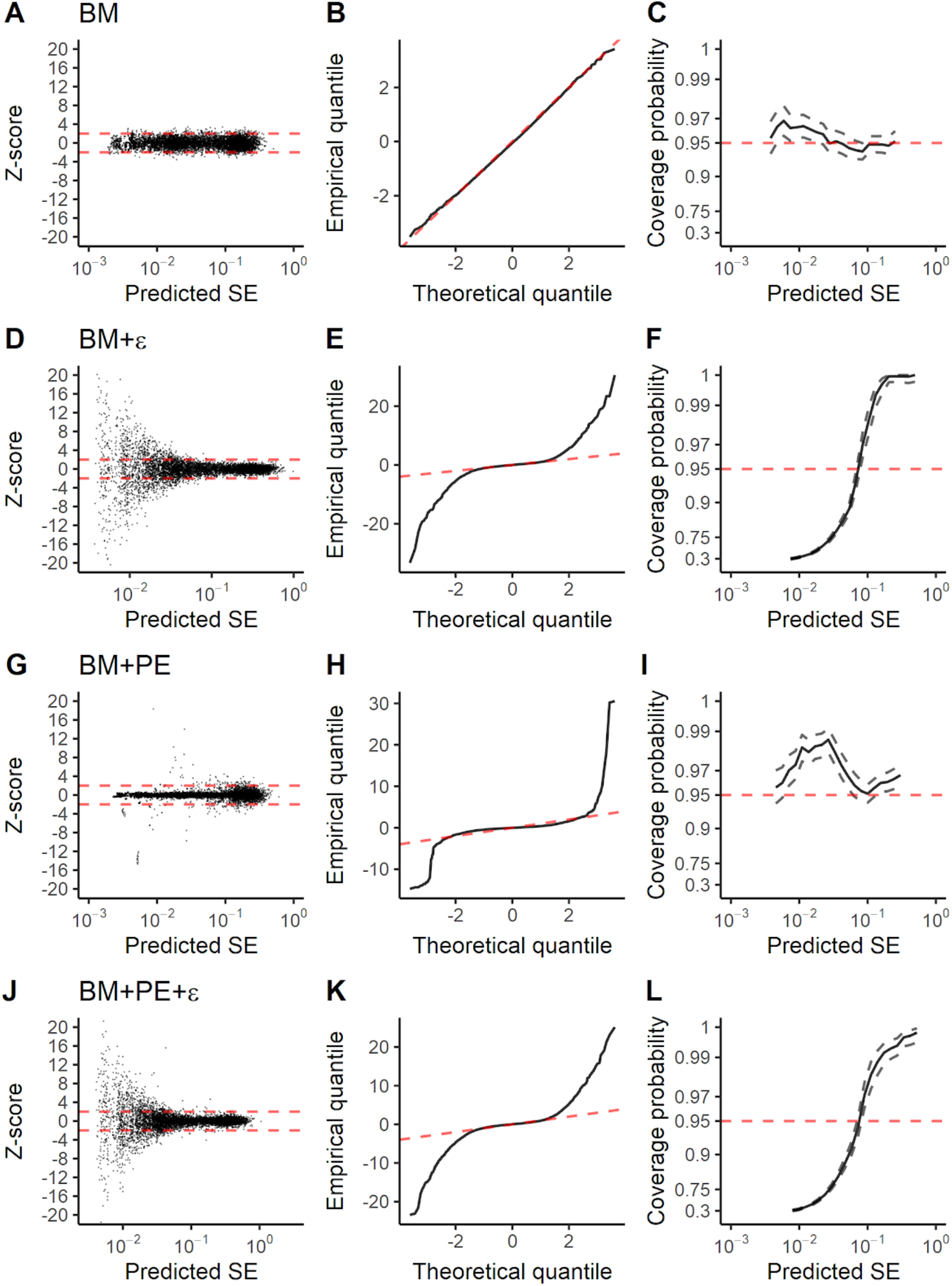
Diagnostic plots of residual errors in ancestral state reconstruction using the BM model on data simulated from various evolutionary models. The layout is the same as that of Figure 1.

**Figure S2.**
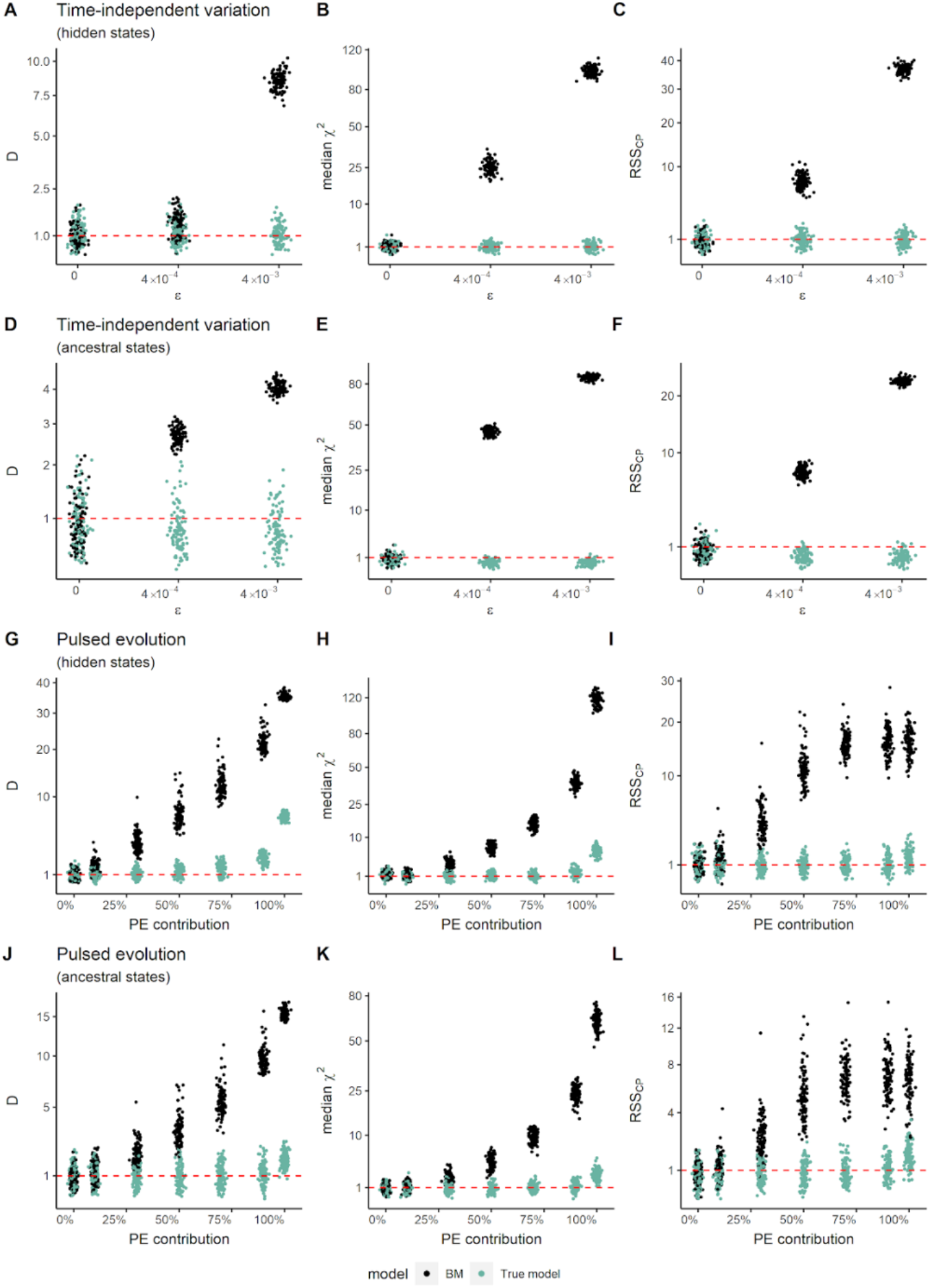
The effect of time-independent variation and pulsed evolution on the reliability of predicted uncertainty. Traits are simulated with varying degrees of time-independent variation and pulsed evolution. We ran 100 replicates for each variation. (A-C) The trend of D statistic, median **χ**^2^ and RSS_CP_ in hidden state prediction in leave-one-out cross-validation when the magnitude of time-independent variation varies. (D-F) The trend of the three statistics in ancestral state reconstruction when the magnitude of time-independent variation varies. (G-I) The trend of the three statistics in hidden state prediction in leave-one-out cross-validation when the contribution of pulsed evolution varies. (J-L) The trend of the three statistics in ancestral state reconstruction when the contribution of pulsed evolution varies. The black and green dots represent the values when the BM model and true model (BM+PE or BM+ε) are used in prediction respectively. The red dashed line in all panels represents the BM baseline. The axes of D statistic, median **χ**^2^ and RSS_CP_ are square-root transformed.

**Figure S3.**
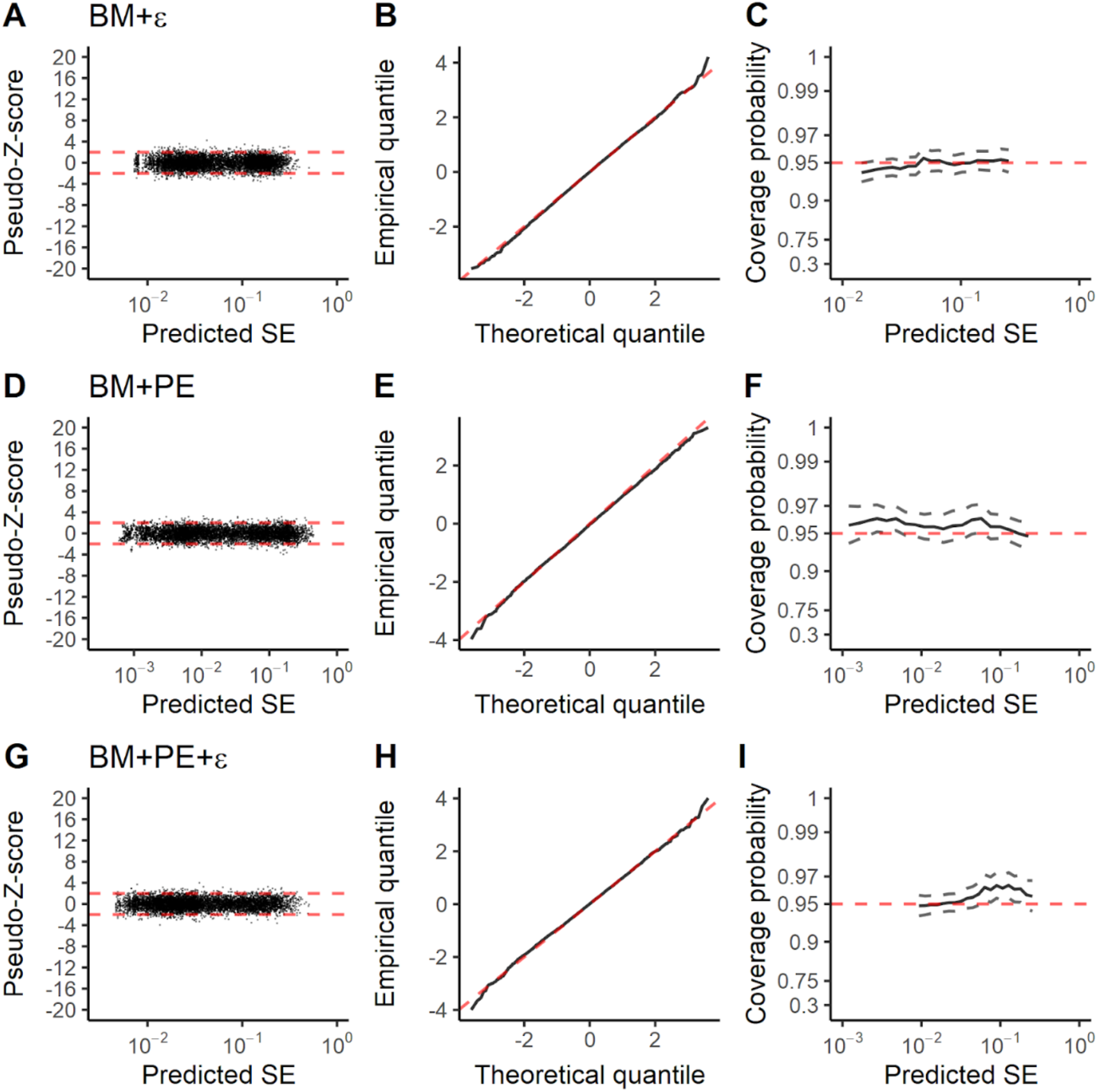
Diagnostic plots of residual errors in ancestral state reconstruction by RasperGade using the true model on data simulated with various evolutionary models. The layout is the same as that of Figure 3.

**Figure S4.**
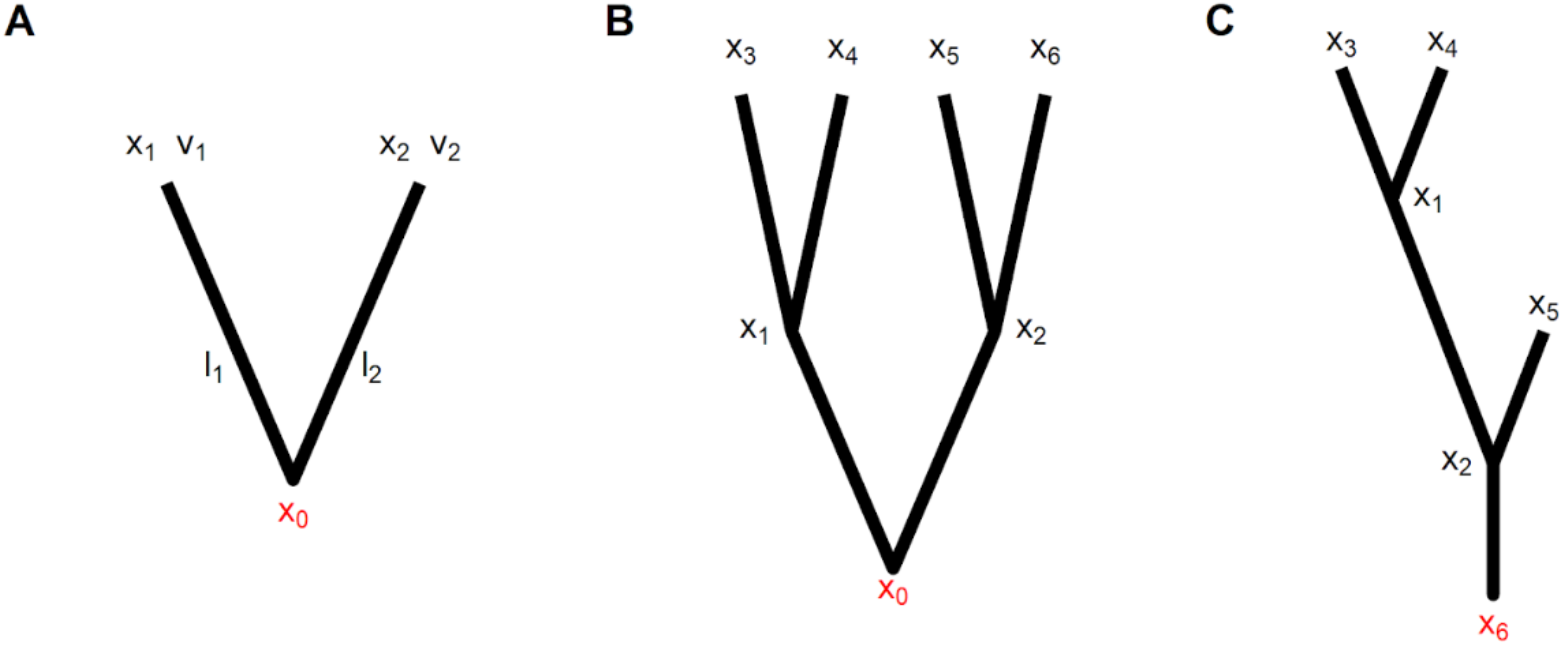
Schematic of ancestral state reconstruction and hidden state prediction. The ancestral or hidden states to be predicted are highlighted in red. *x*: the trait value of a node. *v*: the variance (uncertainty) of the trait value. *l*: the length of a branch. (A) A simple example of ancestral state reconstruction from two descendants. (B) An example of ancestral state reconstruction from tips to the root. (C) An example of hidden state prediction by rerooting the tree with the to-be-predicted node as the root.

**Table S1.** Model testing and the fitted parameters of the BM+PE+ε model.

| Trait | AIC | | Parameters of BM+PE+$\epsilon$ | | | | |
| --- | --- | --- | --- | --- | --- | --- | --- |
| | BM | BM+$\epsilon$ | BM+PE<br>+$\epsilon$ | $\lambda$ | M | r | $\epsilon$ |
| rRNA GC% | -39060 | -39209 | **-42012** | 2.13 | $4.87 \times 10^{-3}$ | $1.84 \times 10^{-3}$ | $1.83 \times 10^{-6}$ |
| Genomic GC% | -28106 | -28104 | **-30590** | 6.30 | $1.42 \times 10^{-2}$ | $8.20 \times 10^{-3}$ | $1.09 \times 10^{-5}$ |
| Genome size | -11009 | -14722 | **-15638** | 7.21 | $2.91 \times 10^{-2}$ | $4.46 \times 10^{-2}$ | $1.29 \times 10^{-3}$ |
| N-ARSC | -35506 | -40743 | **-41703** | 17.30 | $2.09 \times 10^{-4}$ | $1.91 \times 10^{-4}$ | $3.00 \times 10^{-5}$ |
| 16S rRNA GCN | 28182 | 19305 | **15721** | 8.48 | 1.23 | $3.60 \times 10^{-7}$ | $2.61 \times 10^{-2}$ |
| Body size | 8607 | 2068 | **1407** | $3.10 \times 10^{-2}$ | $1.25 \times 10^{-1}$ | $2.14 \times 10^{-9}$ | $2.52 \times 10^{-2}$ |

Note: The table presents the AIC of fitted models and the parameters of the best model. AIC: Akaike Information Criterion of each fitted model. $\lambda$ : The number of jumps per unit branch length. M: The variance of trait change per jump. r: the rate of BM. $\epsilon$ : the variance of time-independent variation.

**Table S2.** The performance of ancestral state reconstruction using the BM model and the true model.

| Model used in simulation | Model used in ancestral state prediction | Performance statistics |  |  |  |  |
| --- | --- | --- | --- | --- | --- | --- |
| | | Error | Bias | D | Median $\chi^2$ | RSS<sub>CP</sub> |
| BM | BM (baseline) | $6.316 \times 10^{-2}$ | $-3.98 \times 10^{-4}$ | 1.00 | 1.00 | 1.00 |
| BM+$\epsilon$ | BM | $8.196 \times 10^{-2}$ | $-5.18 \times 10^{-4}$ | 4.02*** | 85.29*** | 23.19*** |
| | BM+$\epsilon$ | $8.142 \times 10^{-2}$ *** | $-5.36 \times 10^{-4}$ | 0.89 | 0.59 | 0.59 |
| BM+PE | BM | $5.508 \times 10^{-2}$ | $-3.67 \times 10^{-4}$ | 9.81*** | 23.79*** | 6.84*** |
| | BM+PE | $4.227 \times 10^{-2}$ *** | $-2.19 \times 10^{-4}$ | 1.15** | 1.12* | 1.08 |
| BM+PE+$\epsilon$ | BM | $7.751 \times 10^{-2}$ | $2.72 \times 10^{-4}$ | 4.08*** | 86.20*** | 19.04*** |
| | BM+PE+$\epsilon$ | $6.601 \times 10^{-2}$ *** | $1.49 \times 10^{-4}$ | 0.91 | 0.66 | 0.68 |

Note: same as the note in Table 1.

